# Online control of prehension predicts performance on a standardised motor assessment test in 8-12 year-old children

**DOI:** 10.1101/095703

**Authors:** Caroline C.V. Blanchard, Hannah L. McGlashan, Blandine French, Rachel J. Sperring, Bianca Petrocochino, Nicholas P. Holmes

## Abstract

Goal-directed hand movements are guided by sensory information and may be adjusted ‘online’, during the movement. If the target of a movement unexpectedly changes position, trajectory corrections can be initiated in as little as 100ms in adults. This rapid visual online control is impaired in children with developmental coordination disorder (DCD), and potentially in other neurodevelopmental conditions. We investigated the visual control of hand movements in children in a ‘centre-out’ double-step reaching and grasping task, and examined how parameters of this visuomotor control co-vary with performance on standardised motor tests often used with typically and atypically developing children. Two groups of children aged 8-12 years were asked to reach and grasp an illuminated central ball on a vertically oriented board. On a proportion of trials, and at movement onset, the illumination switched unpredictably to one of four other balls in a centre-out configuration (left, right, up, or down). When the target moved, all but one of the children were able to correct their movements before reaching the initial target, at least on some trials, but the latencies to initiate these corrections were longer than those typically reported in the adult literature, ranging from 211 to 581 ms. These later corrections may be due to less developed motor skills in children, or to the increased cognitive and biomechanical complexity of switching movements in four directions. In the first group (n=187), reaching and grasping parameters significantly predicted standardised movement scores on the MABC-2, most strongly for the aiming and catching component. In the second group (n=85), these same parameters did not significantly predict scores on the DCDQ-07 parent questionnaire. Our reaching and grasping task provides a sensitive and continuous measure of movement skill that predicts scores on standardized movement tasks used to screen for DCD.

## Introduction

Almost from the moment able-bodied people wake up, they begin reaching and grasping for objects with their hands – bed covers, a cup of coffee, a toothbrush. Coordinating and controlling accurate, goal-directed reaching and grasping movements is done many times a day. Visually-guided movements longer than about 100ms in duration may benefit from visual online control (Castiello et al., 1991; Farnè et al., 2003; Paulignan et al., 1991a,b; Tresilian, 2012) which is the ability to quickly and accurately correct one’s movement in response to unexpected changes in the hand or target’s position or orientation, for example, when grasping an object as it is falling from your desk (Ruddock et al., 2014). In such situations, the reaching movement must be altered online, to reduce the error and bring the hand and target closer together. This online error correction occurs for many goal-directed movements, but takes some time. The most rapid movement corrections in adult humans begin at 90-120ms after an unexpected change in target object position (Paulignan et al., 1991a); the movement towards the initial target must be cancelled, and an acceleration towards the new target must be programmed. Adjustments to the reaching component of prehension (i.e., hand position) based on changes in object position occur more rapidly than adjustments to the grasping component (hand orientation and grip aperture) based on changes in object size (Paulignan et al., 1991a,b).

Visual online control is an important part of theories of motor control in which limb movements are controlled by internal feedback loops, which are continuously updated to adjust for error and changes in the environment (Goodale et al., 1986; Hyde and Wilson, 2011a, 2011b, Paulignan et al., 1991a, 1991b; Prablanc and Martin, 1992; Wilson et al., 2013). The feedback loops integrate sensory input and motor output to adjust the ongoing motor commands. A review of internal feedback models suggests that accurate arm movements cannot be executed purely under feedback control because visual feedback loops are too slow (Wolpert et al., 1998). Instead, internal models of the body in the brain allow for ‘forward’ predictions of the likely sensory consequences of ongoing actions so that these likely consequences can be taken into account when correcting movements, in advance of actual feedback.

In experimental settings, online movement corrections can be studied using a ‘double-step’ perturbation task, which involves the participant rapidly changing their movement from one target towards another target location (after a ‘perturbation’ of the target position) before the initial movement is complete (Hyde and Wilson, 2011a; Paulignan et al., 1991a, 1991b; Prablanc and Martin, 1992; Van Braeckel et al., 2007). Wilson and Hyde (2013) used a double-step reaching task to explore age-related changes in visual online control in children. They found that older and mid-aged typically developing (TD) children corrected their reaching during the perturbed trials of the task significantly faster than younger children. They also found that adults were faster than older children on all measures.

This double-step reaching experimental paradigm has also been used to explore visual online control in children with developmental coordination disorder (DCD, Hyde and Wilson, 2011a, 2011b, 2013; Plumb et al., 2008). DCD, sometimes referred to as developmental dyspraxia, or just dyspraxia, is a complex neurodevelopmental disorder and has a prevalence of around 6% in school-age children (American Psychiatric Association, 2000). The DSM-5 diagnostic criteria for DCD includes the disturbances in acquisition and execution of basic motor skills, to the extent that it interferes with daily activities and impacts the child’s life both at school and during their leisure time, with an early onset during the developmental period, and that can’t be better explained by any other disability (American Psychiatric Association, 2013). Plumb and colleagues (2008) conducted the first study exploring visual online control in children with DCD and did not find evidence for children with DCD having a specific disruption in this domain. However, the authors cautiously noted that performance in their sample was globally so poor that it was not possible to determine where the deficit lay. Instead they supported a more fundamental movement dysfunction that makes it very difficult to pinpoint a specific mechanism. In Plumb and colleagues’ study, children stood up and made an aiming movement using a stylus towards a target which changed location unexpectedly on some trials. As they had difficulty performing the task standing, children with DCD were allowed to sit down during the task, and the hand-held stylus was made thicker for them than for the TD children. Plumb and colleagues’ results showed that children with DCD took longer to complete the task overall, but there was no significant interaction between condition (perturbation versus non-perturbation) and group (DCD versus TD). As the authors stated, the ability to adjust to perturbations might be related to the quality of motor commands and/or to the quality of the feedback controller. Thus, observing difficulties with visual online control doesn’t necessarily imply problem entirely at the level of the feedback controller. However, since the procedure was different for the TD children and children with DCD, the absence of evidence for specific deficits in visual online control was later re-assessed (Hyde and Wilson, 2011a).

Evidence in support of specific deficits in online control in children with DCD comes from later studies (Hyde and Wilson, 2011a, 2011b; Wilson and Hyde, 2013). Hyde and Wilson (2011a) used a computerised visual online control task, with targets displayed on a LCD touch-screen. Children had electromagnetic sensors attached to their index finger, via a glove, that recorded its position. The authors found that children with DCD displayed longer movement times and increased error rates when responding to target perturbations during the visual online control task. They also found that the performance of children with DCD aged eight to twelve years old was equal to that of typically developing five to seven-year-old children, in regards to rapid online control (Wilson and Hyde, 2013).

The foregoing work on online control has compared groups of children with DCD to TD children, but has not examined how children’s visual online control across a wide range of movement skills covaries with performance on the standardised tests of movement coordination. By testing children both with and without motor impairments, and by assessing a wide range of movement variables on a continuous scale, the present study explores which reaching and grasping parameters best predict scores on standardised measures of movement ability often used for assessing children with DCD - the Movement Assessment Battery for Children 2^nd^ Edition (MABC-2; Henderson, S.E. et al., 2007), and the Developmental Coordination Disorder Questionnaire (DCDQ’07; Wilson et al., 2000, 2009).

As well as assessing performance across a continuous scale of movement skill, our work is based on a double-step reaching-and-grasping task, involving four alternative possible movement directions, in contrast to the typical two alternative targets used in many prior studies (though see, e.g., Prablanc and Martin, 1992). This more unpredictable displacement of the target object is more like a real-world problem, and reduces both the potential overlearning of a small number of target locations, and the usefulness of movement strategies such as ‘reach midway between the targets, then wait to see if anything changes’. Further, instead of presenting targets on a flat, 2D computer screen, which may result in motion blur and a lack of reliable and precise onsets and offsets of the displayed stimuli (Elze and Tanner, 2012), we used LEDs to illuminate, with millisecond precision, translucent table tennis ball targets that were physically grasped by the children. Our aim here is to examine in detail the relationships between visual online control and standardised movement scores in children aged 8-12 years, across a wide range of movement coordination skill.

## Material and Methods

### Participants

A total of 299 children were studied. After removal of 48 datasets because of electromagnetic artefacts and other outliers (Figure 2), 187 children performed the reaching and grasping task and the MABC-2 (109 females, mean ± SD age = 9.30 ± 0.74 years), and 85 children (46 females, mean ± SD age = 10.34 ± 0.95 years) participated as part of Nottingham University’s public Summer Scientist Week, 2015. For these children, the parents had completed the DCDQ’07 questionnaire. The children were recruited in different ways, including through their teachers, using a local database of schools, directly through parents or caregivers using a local database of individual participants, or by other means (e.g., by their expressing an interest directly or by email during or after outreach work). All the children had normal or corrected vision. All parents and children gave written, informed consent and assent, respectively. The experimental procedures were approved by the local ethical review committees at the Universities of Nottingham and Reading, and were in accordance with the Declaration of Helsinki (as of 2008).

**Table 1:** Factors extracted from 87 kinematic variables, and their relationship with MABC-2 and DCDQ-07 scores

| Factor | Loadings<br>≥±0.3 (N) | Description† | Variance<br>explained (%) | Correlation with MABC-2 component scores, $r_{185}$<br>(stepwise regression coefficient±SE) | | | | Correlation with<br>DCDQ-07 scores, $r_{83}$ | | | |
| --- | --- | --- | --- | --- | --- | --- | --- | --- | --- | --- | --- |
|  |  |  |  | MD | A&C | B | T | C | F | G | T |
| 1 | 50 | - | 25.4 | .050 | .020 | .051 | .054 | .019 | .111 | .069 | .069 |
| 2 | 40 | - | 13.0 | -.040 | .010 | -.016 | -.023 | -.127 | -.068 | -.068 | -.106 |
| 3 | 31 | - | 10.3 | .022 | -.067 | -.007 | -.014 | -.086 | -.024 | -.048 | -.066 |
| 4 | 18 | 'Temporal variability; slow<br>corrections' | 6.48 | .050 | .134 | .111 | .118 | -.068 | .118 | .032 | .016 |
| 5 | 18 | - | 5.24 | -.186* | -.140 | -.183* | -.217* | -.202 | -.016 | .011 | -.093 |
| 6 | 14 | 'Jerk, path, correction<br>latency, curvature' | 4.48 | -.210* | -.357* | -.182* | -.294* | -.079 | -.123 | -.051 | -.093 |
|  |  |  |  | (-1.55±0.537) | (-1.77±0.344) |  | (-4.77±1.14) |  |  |  |  |
| 7 | 8 | 'Additional movement during<br>corrections' | 2.99 | .031 | -.139 | .169 | .055 | .223 | .110 | .130 | .187 |
| 8 | 3 | 'Early peak grip aperture' | 2.51 | .000 | -.064 | -.034 | -.036 | .067 | -.090 | .038 | .020 |
| 9 | 5 | 'High correction magnitude<br>variability; long paths' | 2.45 | .089 | .060 | .032 | .074 | -.194 | -.038 | -.143 | -.158 |
| 10 | 4 | 'Additional acceleration/jerk;<br>correction variability | 2.12 | .020 | -.031 | .027 | .013 | .037 | .085 | -.050 | .022 |
| 11 | 5 | 'Low correction magnitude<br>and grip timing variability' | 2.06 | .129 | .177 | .103 | .163 | .038 | -.146 | .002 | -.024 |
| 12 | 4 | 'Smaller, later peak grip<br>aperture' | 1.76 | .044 | .032 | .091 | .074 | .034 | .132 | .122 | .103 |
| 13 | 5 | 'Slow RT; long correction<br>paths' | 1.66 | -.047 | .036 | .129 | .053 | -.122 | -.057 | -.131 | -.125 |
| 14 | 0 | ('Path variability') | 1.53 | .213* | .136 | .148 | .211* | -.036 | -.001 | -.049 | -.037 |
| 15 | 3 | 'Variable and late peak grip<br>aperture' | 1.32 | .235* | .194* | .199* | .264* | .196 | .231 | .215 | .244 |
|  |  |  |  | (1.68±0.517) | (0.918±0.332) | (1.54±0.573) | (4.12±1.10) |  |  |  |  |
| 16 | 2 | 'Long reaction time' | 1.29 | -.134 | -.092 | -.082 | -.129 | -.092 | -.148 | -.054 | -.107 |
| 17 | 2 | 'Variable peak grip aperture;<br>straight reaches' | 1.16 | -.001 | -.164 | -.083 | -.091 | -.091 | -.130 | -.118 | -.127 |

† Descriptions are subjective and approximate; a full list of factor coefficients is provided in the Supplementary Tables. MD: Manual dexterity;
A&C: Aiming and catching; B: Balance; T; Total; C: Coordination during movement; F: Fine motor; G: General. \*p≤.0125 (corrected for 4
comparisons per factor). Values in parentheses are the coefficients ± standard errors of the factors that remained as significant predictors in a stepwise linear regression ( $p < .0125$ ).

### Motor and cognitive skills assessment procedures

#### Movement Assessment Battery for Children, 2^nd^ Edition

(Henderson et al., 2007) – The MABC-2 is a standardised test used to assess motor coordination impairments in children and adolescents. Children directly perform a set of eight tasks among three components, with three tasks assessing manual dexterity, two assessing aiming and catching, and three assessing balance. Although the skills tested are the same for all, different tasks are designed for three specific age bands: age band 1 (3-6 years), age band 2 (7-10 years) and age band 3 (11-16 years). Each raw score obtained by a child in each of the eight tasks is then converted into an item standard score following a scoring table depending on the child’s age within the age band (i.e., for age band 2, 7:0-7:11, 8:0-8:11, 9:0-9:11, 10:0-10:11). These scores are summed into a component score, then converted into standard scores (mean=10, SD=3) with their equivalent percentiles for the three component scores and the total of the MABC-2. To facilitate calculation of standard and component scores, we created macros in Excel to automate this process by extracting the appropriate scores from look-up tables (Supplementary data). In the current study, the tasks were age-appropriate, with all children performing tasks from the 7-10 year old bracket (several children over 10 years of age were tested only with the DCDQ'07, and not with the MABC-2).

#### DCDQ’07

(Wilson et al., 2000, 2009) – The DCDQ’07 is a brief parent questionnaire designed to screen for motor problems associated with DCD in children aged 5 to 15 years. Parents are asked to compare their child’s motor performance to that of his/her peers depending on the child age band (5:0-7:11, 8:0-9:11, 10:0-15:0). The DCDQ’07 consists of 15 items grouped into 3 areas: control during movement, fine motor/handwriting, and general coordination. For children aged 8 to 10 years, a score of 15-55 suggests the kinds of motor problems associated with DCD, whereas a score of > 55 probably does not indicate such problems. For children aged 10 to 15 years, a score of 15-57 suggests motor problems associated with DCD, where as a score of > 57 probably does not.

#### Online control measure

We used a centre-out double-step reaching and grasping task to measure how children alter their movement when reaching to grasp an illuminated ball with their dominant hand. Children were seated comfortably at a table with a vertical (40×50 cm) board on the table 30 cm in front of their hand, which was placed on a starting position, 30 cm from the board. Five translucent orange table tennis balls (4 cm diameter) were attached at the centre, top, bottom, right and left sides of the board, with the centres of the four eccentric balls 11.5 cm away from the centre (Figure 1).

**Figure 1:**
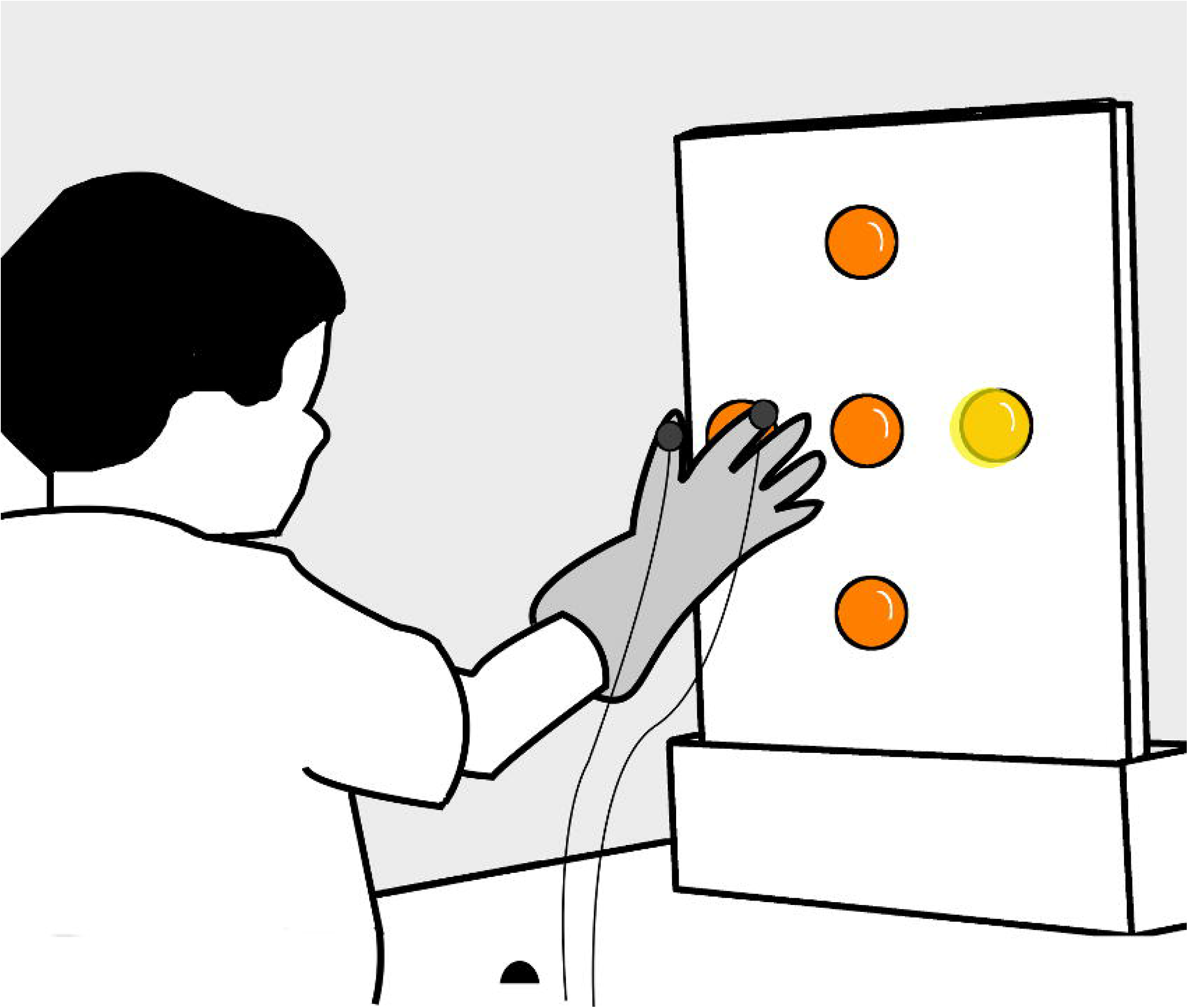
Experimental apparatus. Children sat and rested their hand on a table, keeping their index finger and fingers in a ‘start’ location (black hemisphere), 30cm away from a vertical board with five orange table tennis balls attached at the centre, top, bottom, right and left sides - the centres of the four eccentric balls were 11.5cm away from the centre. Children wore a glove with trackers (solid grey circles) attached on the index finger and thumb.

At the start of each trial, after a brief interval (randomly 1-3 s) an ultra-bright white LED illuminated the single central target ball from the inside. On 60% of the trials for the first 48 children, and 50% of trials for the last 251 children, the central ball remained lit, but on the remaining 40% (50%) of trials at the onset of the participant’s hand movement the illumination switched from one ball to another in a centre-out configuration. We changed the proportion of trials after the first 48 children in order to have one additional (critical) trial per eccentric ball position. While many researchers present 80% ‘unperturbed’ and 20% ‘perturbed’ target conditions, this distribution is not universal (e.g., Prablanc and Martin, 1992, used 33% unperturbed and 67% perturbed), and neither perturbation probability nor perturbation expectation have strong effects on the latency to initiate movement corrections (Cameron et al., 2013). The hand position was analysed online, and the target change occurred as soon as the tangential velocity of the thumb reached 15 cm/s. This criterion was subsequently changed (after the first 48 children), to a velocity towards the central target of 15 cm/s in order to counter the strategy of some children opting to make a short initial movement (e.g., upwards or sideways), before making a second movement towards the target location, which may since have changed. The balls in the up, down, left, and right positions each lit up on 10% (12.5%) of the trials in a pseudorandom order. Children were instructed to start each trial with their thumb and index fingers closed in a pincer grip and placed on the starting point, then to reach and grasp the illuminated ball as accurately and as quickly as possible, but in a controlled manner – as natural a reaching movement as possible. Children were instructed to interrupt their movement to the central ball and grasp instead the eccentric ball when the illumination switched locations. There were 10 practice trials before the main data collection to familiarise the children with the task and this was followed by one testing block of 40 trials. Motion trackers were attached over the thumb and index fingers (i.e., the grasp ‘opposition axis’, Holt et al., 2013) of a ‘NASA’ astronaut’s glove (this did not appear to affect children’s hand movements, see also <u>Hyde and Wilson, 2011b)</u>, to record the position (3 degrees of freedom) of these digits with a Polhemus Fastrack (Polhemus, Colchester, VT, USA) magnetic tracking system. The system has a spatial accuracy of 0.08 cm, and a precision of 0.0055 cm (for the average location sampled in the current study), sampling the two receivers, each at 60 Hz. We opted for two trackers sampling at 60Hz as the ideal trade-off between trackers (1-4) and frequency (120-30Hz) - an additional third tracker on the wrist would have entailed a reduction of sampling frequency to 40Hz. Since human hand and finger movements cannot move or oscillate at much more than 30 Hz (Raethjen et al., 2000), and the visual online control of movement takes a minimum of 100 ms, 60 Hz sampling is more than adequate to capture the relevant information required to test our hypotheses.

#### Cognitive assessments

Children in the first, MABC-2 (Age band 2, 7-10 years old), group were assessed with the Reading, Verbal Similarities, and Matrices tests of the British Ability Scales (BAS) (Elliot, 1996), and the Conners 3-AI (Conners, 2008). Children in the second, DCDQ’07, group were assessed with the British Picture Vocabulary Scale (Dunn et al., 1997) and the Strengths and Weaknesses of Attention-Deficit/Hyperactivity Disorder Symptoms and Normal Behaviour Scale (SWAN) (Swanson et al., 2001, 2012). Socioeconomic status was estimated from children’s home postcodes using the UK Government’s Indices of Deprivation (IDACI/IMD).

### Design

All children were assessed with the visual online control test. 187 children were assessed with the MABC-2 (Age band 2, 7-10 years old), and 85 children were assessed with the DCDQ’07. Our study was an exploration to investigate correlations between kinematic variables extracted from reaching and grasping movements, and a) MABC-2 component (manual dexterity, aiming and catching, and balance) and total scores, and b) DDCQ’07 scores (coordination; fine motor; general).

**Figure 2:**
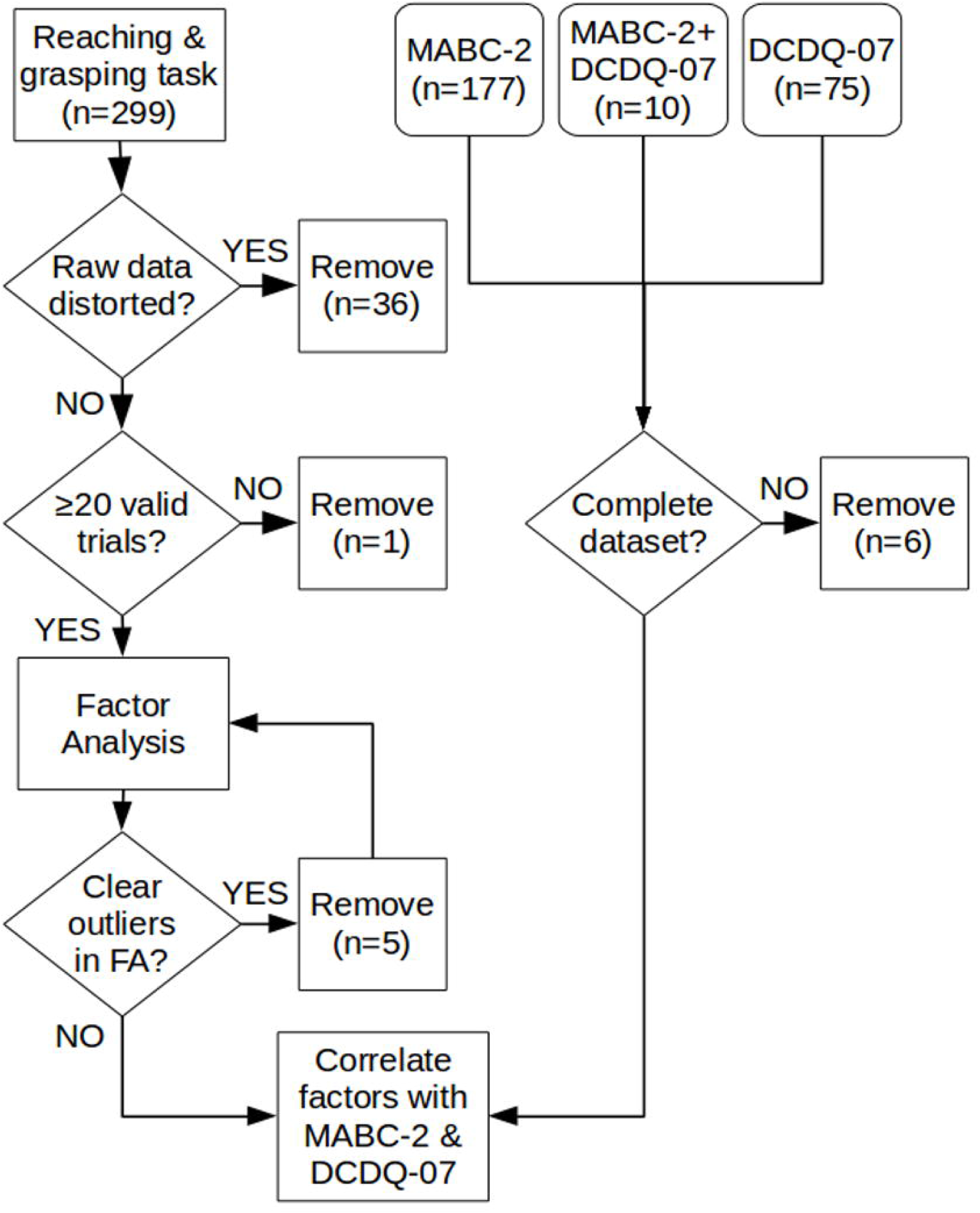
Recruitment and selection. 299 participants were recruited through schools, parents, a DCD support group and a University public engagement event. After removing outliers and artefacts, 262 datasets entered the factor analysis. 181 valid datasets with MABC-2 scores, and 85 valid datasets with DCDQ’07 scores were available for regression analysis. FA: Factor analysis; MABC-2: Movement Assessment Battery for Children-2; DCDQ’07: Developmental Coordination Disorder Questionnaire'07.

### Data analysis

The experiments were run and the data analysis was performed using Matlab (Mathworks, Natick, USA), and SPSS for the factor analysis. All the programs and all raw data are or will be freely available from the last author or his website (Correspondence:http://neurobiography.info/). All data analysis was fully automated and scripted, using procedures developed during previous work (for full methods and discussion, see Holmes and Dakwar, 2015). A summary of the analytical approach is provided here.

#### Raw data

Six degree-of-freedom position and orientation data from the index finger and thumb were acquired at 60 Hz for 2 s per trial. Data were re-sampled to 120 Hz then filtered with a 2^nd^ order, zero-lag (dual-pass) Butterworth filter with a 15 Hz low-pass cut-off. Two raw data channels (index finger and thumb), as well as the mean (used for many kinematic parameters), and the difference (used for grip aperture measures), were processed individually by the same analysis script (hl_kinematics.m, part of the HandLabToolBox). The script is fully-automated, and extracts, from each trial, reaction time (RT, the first sample after 100 ms that exceeds 15 cm/s velocity towards the initial target) and movement time (MT, the first sample after RT in which tangential velocity subsequently remains below 10 cm/s for at least 50 ms; this is combined with target position information, to check whether MT was reached within, or away from the target location, with a 6 cm tolerance), along with peak acceleration (PA, and the time that PA was reached, TPA), peak velocity (PV, TPV), peak deceleration (PD, TPD), path length, mean velocity (MV), movement symmetry (TPD/MT), movement shape (PV/MV), movement curvature (the maximum deviation orthogonal to the straight line joining the locations at RT and MT, divided by the length of that straight line), and root-mean-squared jerk (3^rd^ differential of position over time) and snap (4^th^ differential). All temporal parameters (TPA, TPV, TPD, MT) are expressed relative to movement onset (i.e., after subtracting RT from the time since the target appeared). The difference between index and thumb positions (i.e., grip aperture) was analysed similarly, yielding measures of peak grip aperture (PGA, TPGA).

#### Processed data

The analysis routines then processed the data from each trial of each participant, rejected artefacts, determined errors and outliers on a number of criteria. Exclusion criteria were set after an initial analysis, examining the histograms of extracted parameters, and setting limits to exclude only clear outliers during a second analysis. These ‘outliers’ were all caused by participant error (e.g., moving before target onset, or failing to move), or by hardware failure (e.g., the eccentric target light failing to illuminate, magnetic distortion or interruption of the tracker signal). Details of the trials removed are provided in supplementary data. All subsequent calculations were performed on valid trials only, then were summarised per condition (target location) and participant. Where possible, data were extracted from individual trials. Thus, for each participant, condition, and parameter, mean, SD, and N are available. Temporal parameters (TPA, TPV, TPD, TPGA) were also expressed relative to total MT (TPA/MT, TPV/MT, TPD/MT, TPGA/MT). Summary descriptive statistics are provided for all variables in supplementary data. All raw and summary data were inspected visually, in order to set criteria and adjust analytic parameters and procedures. The final analysis is fully-automated and repeatable. The only human intervention in the final analysis was to exclude two clear outlying participants, following plotting of the factor analysis data – factor analysis is sensitive to outliers (Flora et al., 2012).

#### Correction movements

The principal variables of interest were the latencies, velocities, and accelerations of the corrections made to the reaching component of the movement following target perturbations. By ‘correction movements’, we mean the velocity of the hand towards the (new, perturbed) target location on trials with a change in target location, minus the same component of velocity on trials without a change in target location. This can be measured in several ways. Following previous work (Holmes and Dakwar, 2015; Oostwoud Wijdenes et al., 2014; Veerman et al., 2008), we used the optimal method (Holmes and Dakwar, 2015) – extrapolating back from the peak correction velocity to the start of the correction velocity curve for each trial (Figure 3). The zero-crossing point on the x-axis is found by extrapolating back from the line joining the 25% and 75% points, relative to the maximum correction velocity (Veerman et al., 2008). This was done both on individual trials as well as on the mean trajectories from trials of the same condition (i.e., right, lower, left, or upper targets), and was implemented by a HandLabToolBox function, hl_kinematics_correction.m. To aid comparison with previous similar work, we also calculated the correction time as the ‘additional movement time’ required (Hyde and Wilson, 2011a, 2011b), by subtracting the mean MT on trials without a change in target location from trials with a change.

To visualise the data, the velocities, accelerations, and jerks across trials in the same condition were averaged by aligning the movement onsets. Each trial was also resampled to 120 data points, from RT-5 to MT+5 samples. These resampled, standardised, data were then re-scaled to a maximum height of 1, averaged across conditions per participant, and re-scaled again to a maximum of 1. This re-scaling compensates for between-participant differences in movement velocity, duration, and variability. The final average movement profiles (Figure 4) are then useful to assess the overall ‘quality’ or ‘shape’ of movement.

This analysis revealed a clear progression of velocity and acceleration profiles both as a function of age (Blanchard et al., in preparation), and movement coordination ability (Figure 4). Based on this, we also extracted a number of variables in an attempt to measure the overall shape of movement. The area under the velocity curve between RT and MT is equivalent to the path length (i.e., the integral of velocity over time is distance covered), and similar measures can be extracted for the area under velocity, acceleration, and jerk curves, both for the raw, and the resampled standardised data, for both overall 3D velocity, and the component of velocity in the direction of the target change. In our previous work, we found the component of velocity towards the target provided better measures of movement correction (Holmes & Dakwar, 2015). Finally, the additional velocity, acceleration, and jerk on trials with compared to without a change in target location was calculated.

#### Factor analysis

The typical parameters extracted from position data are highly collinear (Naish et al., 2013). For example, a movement which reaches peak acceleration early will likely also reach peak velocity early; higher acceleration leads to higher velocity; these parameters are correlated. Rather than examine a series of kinematic parameters independently, reducing these highly-correlated variables to a smaller number of more independent factors helps resolve problems with multiple comparisons across different dependent variables. We extracted 87 reaching and grasping parameters from each of 262 participants who had valid reaching and grasping data, and reduced this to 17 factors using principal components analysis in SPSS 21 with oblique (direct oblimin) factor rotation in order to minimise the number of variables loading heavily onto each factor. A criterion of eigenvalues >1 was used for factor selection; factor scores were estimated using Bartlett's method. While researchers may disagree over whether and when to use orthogonal or oblique factor rotation, the underlying mathematics is identical, the total variance explained remains the same, and only with criteria external to the factor analysis itself can the usefulness of any particular rotation method be judged (Comrey & Lee, 1992). We assessed the usefulness of the rotation method and the extracted factors by relating their scores to independent measures of movement coordination.

#### Predicting MABC-2 and DCDQ’07 scores with reaching and grasping factors

The factor scores extracted for each participant and factor were correlated individually with the MABC-2 and DCDQ’07 scores. Further, a stepwise linear regression with all 17 factor scores was run to determine which (if any) of the 17 extracted factors could predict MABC-2 or DCDQ’07 scores.

Unless otherwise stated, an alpha level of .0125 was adopted. Since both the MABC-2 and DCDQ’07 contain four separate scores, this alpha level corrects for four independent comparisons for each standardised test. Means are reported to 3 significant figures, SDs to the same number of decimal places as the means.

## Results

A complete table of descriptive summary statistics, along with all the raw data (i.e., participant means), correlations between variables, and factor analysis results is provided in a supplementary Table, and all the raw data are available freely on request. Here, we summarise only a few pertinent variables. All data are mean±SD unless otherwise stated.

**Figure 3:**
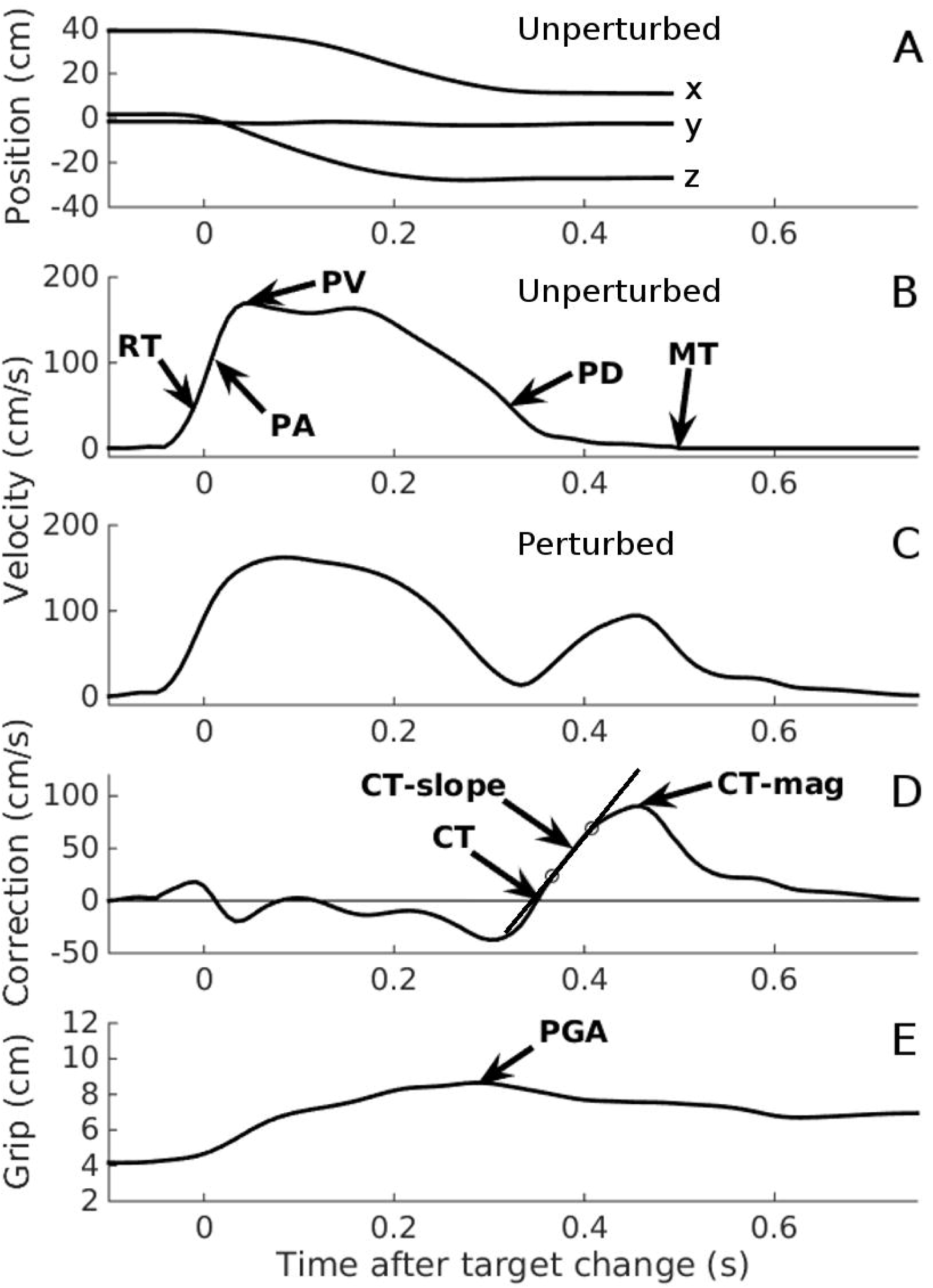
Automated data analysis. Two typical trials, taken from participant number 1, illustrating the automated analysis. Data have been re-aligned to movement onset. **A**. Raw x, y, and z position data from an unperturbed trial. **B**. 3D velocity from the same unperturbed trial indicated in A. RT: Reaction time; PA: Peak acceleration; PV: Peak velocity; PD: Peak deceleration; MT: Movement time. **C**. 3D velocity from a perturbed trial. Note the additional velocity curve, starting at about 0.3 s. **D**. 3D correction velocity: the difference in velocity between the perturbed and an unperturbed trial (for illustration only; the mean unperturbed velocity per participant was used in the analysis). CT: Correction time, determined by extrapolating the line joining the 25% and 75% points on the correction velocity curve (circles), back to the x-axis (Holmes & Dakwar, 2015; Veerman et al., 2008); CT-slope: the slope of the line joining the 25% and 75% lines; CT-mag: the peak correction velocity. **E**. Grip aperture from the same trial as illustrated in C-D. PGA: Peak grip aperture.

### Reaching and grasping task

Overall, 262 children (age=9.65±0.98 years; 154 females; 20 of whom used their left hand for handwriting) completed the reaching and grasping task with sufficient valid trials (≥20, including at least one successful movement correction) for analysis. Of the 40 trials attempted, 32.7±8.8 valid trials per participant were analysed. Across all participants, 52 trials were excluded for not completing within 2 s; 1259 for reaching and stopping on the central target before making a complete movement correction to the correct target (i.e., corrected ‘touchdown’ errors); 35 for central touchdowns which were not followed by a complete movement correction; 196 trials for strong magnetic artefacts or other data corruption; 473 for ‘outliers’ – mostly induced by more subtle magnetic artefacts which were not remedied by initial data filtering. This process led to the removal of 2 participants who no longer had sufficient valid trials for analysis.

Across the valid participants and trials, reaction time (RT) to begin the reaching movement was 364±79 ms (with means ranging between 359 and 371 ms across the five experimental conditions). Movement time (MT) to the central target on unperturbed trials was 514±84 ms; on perturbed trials, movements were completed in means of 731-770 ms. Since a few children did not have valid trials for every target direction, and due to the overall low numbers of perturbed trials, we have not compared movements between the four target locations. There are, however, indications of differences between the left-right and up-down dimensions of movement correction, a finding which will be followed up in a dedicated study.

Using the optimal method from Holmes & Dakwar (2015, Figure 3), the latency to initiate a movement correction following target perturbation was 342±85 ms on average, ranging from 318-335 ms for the right, left, and upper targets, to 390±97 ms for the lower target – this large (≥55 ms) increase in mean latencies for the lower relative to the other targets likely reflects the need to reverse the initial upwards movement required to reach the central target, but this hypothesis needs to be tested with movements starting at the same elevation as the central target. An alternative index of online control measures the additional movement time required to complete the movement correction (MT perturbed minus MT unperturbed, Hyde and Wilson, 2011a). This was 235±90 ms pooled across the four target locations, with right (217±104 ms) and left (214±100 ms) lowest. Unlike for correction latency, additional movement time was greater for the upper (253±101 ms) than the lower (230±103 ms) targets. This difference is discussed below.

To aid data visualisation, tangential 3D velocity profiles were resampled to 120 points between RT-5 samples and MT+5 samples, then re-scaled to a maximum velocity of 1. Standardised velocity profiles were then averaged across trials, conditions, and participants (Figure 4). Visual inspection of these data prompted additional measures of the area under the velocity, acceleration, and jerk curves to be taken. We expected that the apparent between-participant differences in movement shape on perturbed trials might be important predictors of standardised measures of movement coordination.

**Figure 4:**
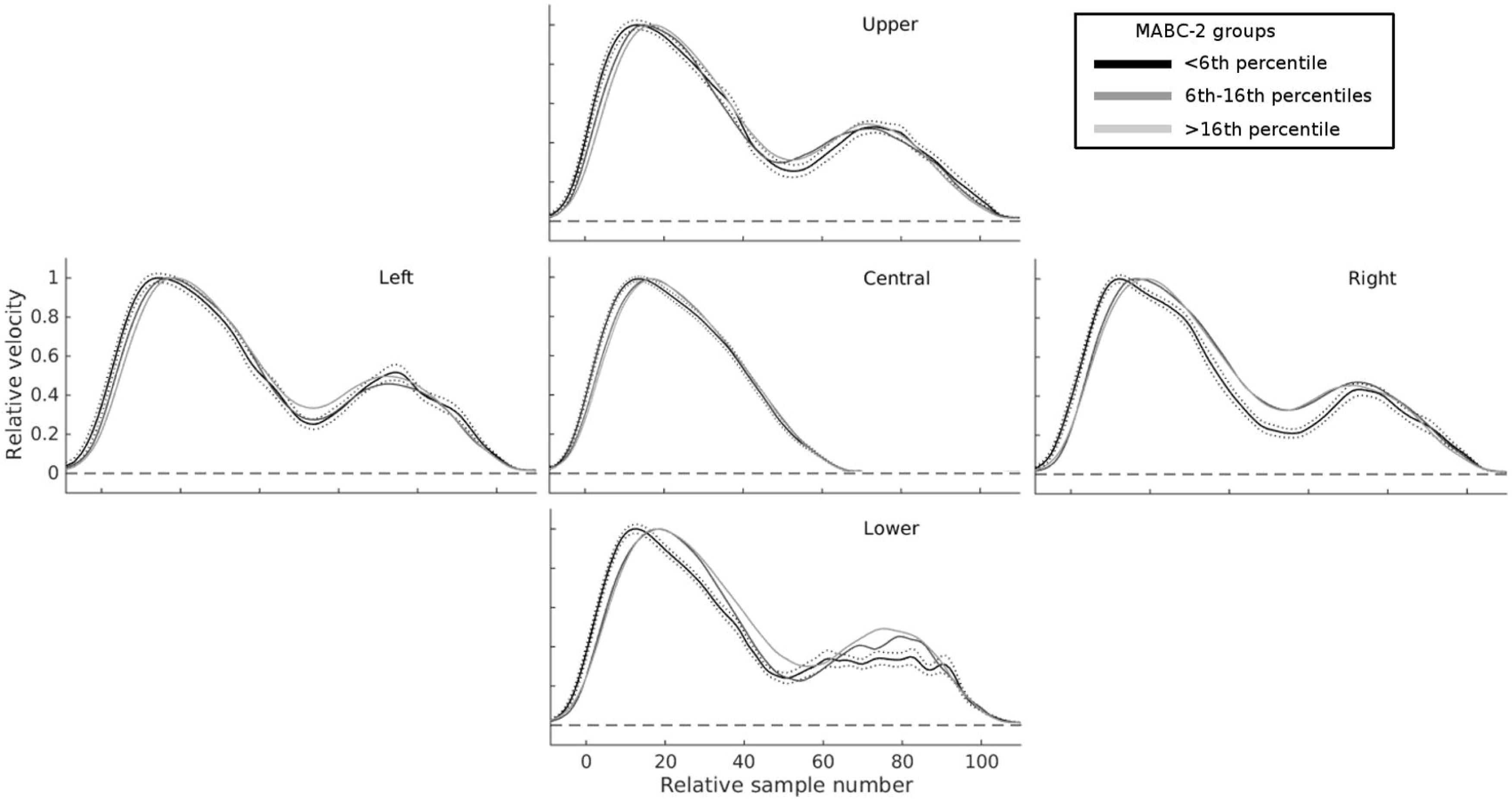
Mean velocity profiles highlight substantial differences in online control as a function of total MABC-2 score. The central panel shows normalised velocity profiles on trials with unperturbed targets, for three groups of children separated according to their overall MABC-2 score: ≤5^th^ percentile (black); >5^th^ and ≤16^th^ percentile (mid-grey); >16^th^ percentile (light grey). The four other panels show velocity profiles for the same three groups in the four conditions with perturbations of target location. For all perturbed targets, the second velocity peak (at around 70-80 samples) is more smoothly integrated with the first in children with higher MABC-2 scores. Children with the lowest scores on the MABC-2 show the largest changes in velocity between the first and second velocity peaks. The broken black lines show the SE for the ≤5^th^ percentile group. The other groups had similar error bars and are not shown for clarity.

### Standardised movement measures

Children assessed with the MABC-2 (n=181, after removing 6 incomplete datasets) achieved a standard score of 9.19±3.40; 25 children were at or below the 5^th^ percentile (i.e., ‘possible DCD’), 35 were between the 6^th^ and 16^th^ percentiles inclusive (i.e., ‘at risk for DCD’, Blank et al., 2012) and 121 were above the 16^th^ percentile. For the DCDQ’07, 85 children’s parents rated them as 61.8±17.9 overall. 27 children had total parent ratings below the cut-off for ‘possible DCD’, and 58 above the cut-off.

### Factor analysis for data reduction

Eighty-seven variables derived from the kinematic data were entered into an exploratory factor analysis with oblique factor rotation. Seventeen resulting factors had eigenvalues >1, accounting for 85.7% of the variance (Table 1). The first three factors had loadings of ≥0.3 on 50, 40, and 31 original variables respectively, and as such were hard to describe, but likely account for general between-participants’ differences in movement speed and variability, or body size, which affects multiple variables. The remaining 14 factors loaded strongly onto between 0 and 18 original variables. An attempt to describe these factors is presented in Table 1, along with the correlation between scores from each factor and the MABC-2 and DCDQ’07 scores. Factors 5, 6, 14, and 15 correlated significantly (p≤0125, 2-tailed) with at least one component of the MABC-2; none correlated significantly with DCDQ’07 scores. The strongest relationship was between factor 6, which explained 4.48% of the reaching and grasping variance, and the MABC-2 aiming and catching component scores (r_179_=-.357, p<.0001). Of the original variables, the strongest relationship between kinematic and standard scores was between movement correction latency measured from individual trials (Holmes & Dakwar, 2015) and the aiming and catching score on the MABC-2 (r_179_=-.260, p=.0004).

### Predicting MABC-2 and DCDQ’07 scores with reaching and grasping factors

Since the factor rotation method was oblique, the resulting factors could still be collinear, however 17 partially collinear factors are more manageable than 87 more highly collinear original variables. Rather than interpret each of the correlations between factors and MABC-2 scores individually, the 17 factor scores were entered as predictors in a stepwise linear regression to identify those factors which explained significant (p-enter≤.0125, p-remove≥.10) variance in the MABC-2 scores. Only two factors, 6 and 15, were retained in the stepwise regression. Factor 6 was the strongest, and 15 the second predictor of both aiming and catching scores, F(2, 173)=17.0, p<.0001, r^2^=0.165, and total MABC-2 scores, F(2, 173)=15.9, p<.0001, r^2^=0.155. Factor 15 was the strongest, and 6 the second, predictor of manual dexterity scores, F(2, 173)=9.48, p=.0001, r^2^=0.099. Finally, factor 15 was the sole predictor of balance scores, F(1, 174)=7.17, p=.008, r^2^=0.040. The regression coefficients are provided in Table 1, and the whole model fits in Figure 5.

The factor analysis and subsequent stepwise regression identified factors 6 and 15 as clear and strong predictors of MABC-2 scores, particularly for the aiming and catching component. Factor 6 loaded most strongly (≥.3) on the additional area under the jerk and acceleration curves in perturbed compared to unperturbed trials, expressed either as a difference (jerk: .512, acceleration: .321) or a ratio (.479, .314), the SD of TPGA (.491), the relative time of peak acceleration (-.420), path length mean (.407) and SD (.354), the SD of MT (.393), the standardised area under the jerk curve (-.372), the mean correction latency per trial (.322), mean curvature (.308), and the SD of jerk (-.302). Since factor 6 was negatively correlated with the MABC-2 scores, larger increases in jerk and acceleration and later corrections on perturbation trials, longer and more curved paths, higher variability of grip timing, MT, and path, along with relatively earlier peak acceleration, and lower jerk predicted lower MABC-2 scores. Factor 15 loaded strongly (≥.3) on the SD of PGA (.380), the relative time of PGA (.377), and the standardised area under the jerk curve (.310). Since factor 15 was positively correlated with the MABC-2, more variable and relatively later peak grip aperture and larger areas under the standardised jerk curves predicted higher MABC-2 scores.

**Figure 5:**
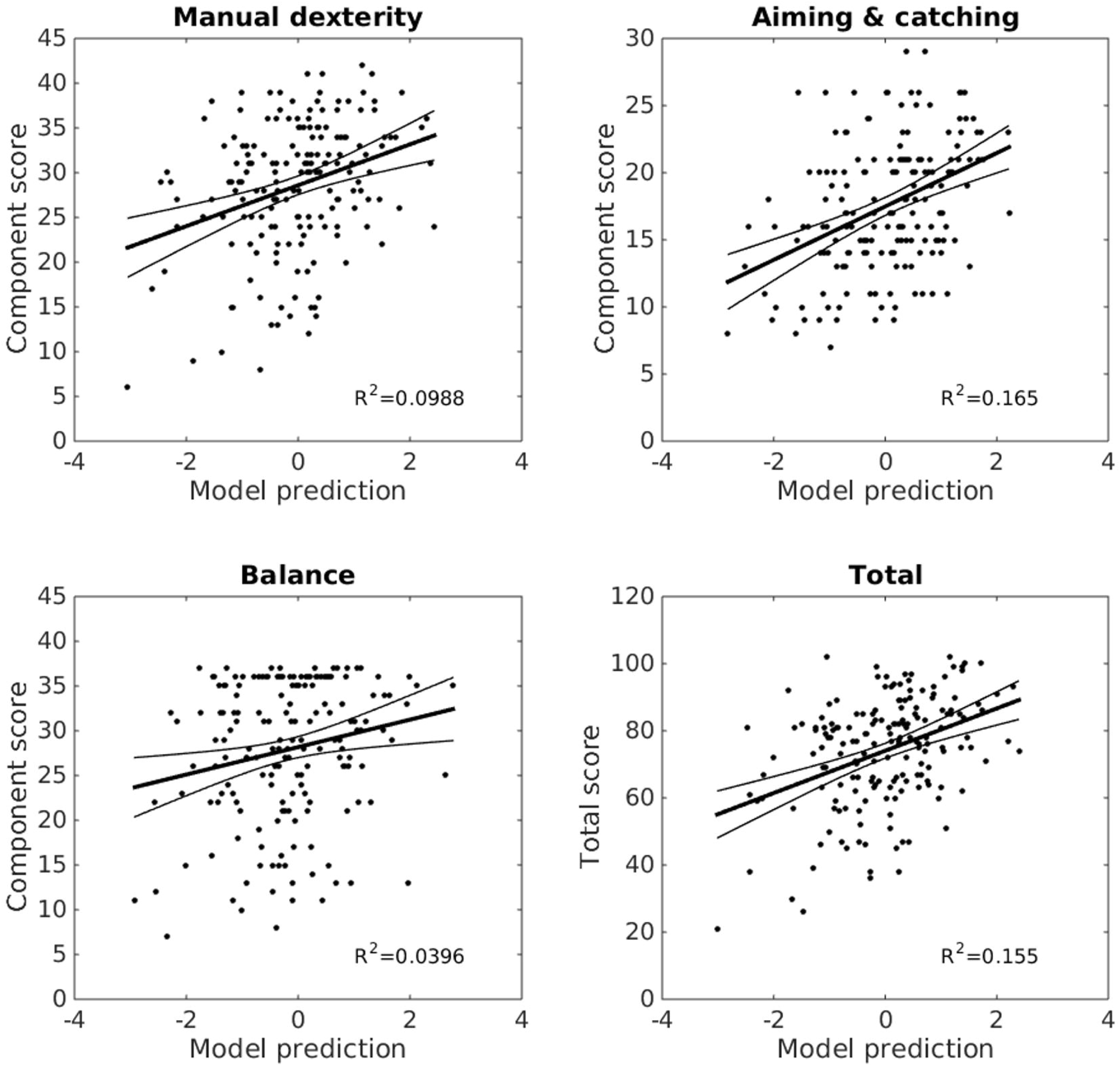
Reaching and grasping factors predict MABC-2 performance. Linear models combining factors 6 and 15 from the factor analysis explain a significant proportion of variance (r^^^2) in children’s MABC-2 performance (y-axes, MABC-2 component and total scores). X-axes show standardised model predictions. The trend line shows the model fit, with 95% confidence intervals. Reaching and grasping explains 17% of the variance in aiming and catching scores, 16% of variance in the total scores, 10% of the manual dexterity, and 4% of the balance score variance.

### Group analyses

The relationships between reaching and grasping and standardised movement coordination scores seem to be continuous, rather than containing any discontinuities at particular scores or ranges. Nevertheless, following a reviewer’s request, the continuum was divided into discrete groups on the basis of both clinical diagnoses (e.g., DCD diagnosis) and the MABC-2 and DCDQ’07 scores relating to clinically significant cut-offs. In our sample, 11 children had formal diagnoses of DCD; 25 children (13.8% of our sample) were at or below the 5^th^ percentile of the MABC-2; 35 (19.3%) were between the 6^th^ and 16^th^ percentile inclusive; and 121 (66.8%) had scores above the 16^th^ percentile. For the DCDQ’07, a large proportion (31.7%) of parents rated their children as having movement coordination below the cut-off. These groups were compared on factors 6 and 15 from the factor analysis, and on the model prediction scores (summary data in the supplementary table).

Children at or below the 5^th^ percentile on the MABC-2 had significantly different (p≤.025, 1-tailed, correcting for 2 comparisons) scores from children above the 16^th^ percentile on both factors 6, t_139_=3.08, p=.001, and 15, t_139_=-2.37, p=.01, and children between the 6^th^ and 16^th^ percentiles inclusive also differed from the >16^th^ percentile group on factor 6 (t_151_=2.19, p=.015), but not factor 15, t_151_=-1.25, p=. 11. Factor scores of the groups at or below the 5^th^ and between the 6^th^ and 16^th^ inclusive did not differ significantly. Regarding the linear model predictions of the MABC-2 component scores and total scores, the same pattern was found, with the two lower-scoring MABC-2 groups differing significantly (p≤.0125, 1-tailed, correcting for 4 comparisons) from the higher-scoring group on manual dexterity, aiming and catching, and total scores, while only the comparison between the ≤5^th^ percentile group and the >16^th^ percentile group was significant for the balance scores. All the differences were in the expected directions, which is not surprising as the models were set up to predict these scores – dividing the range into bins and re-testing is a statistical ‘double-dip’. There was no evidence for significant differences between the 11 children with a formal diagnosis of DCD and the rest of the sample, either on factor 6, t_255_=0.663, p=.51 or factor 15, t_255_=-1.26, p=.21, or on the aiming and catching, balance, or total scores (|t_174_|s<1.68, ps>.095. Again, this may not be surprising, as the model was set up to predict MABC-2 scores, rather than DCD diagnosis. Factor 12, however, did show a relatively large difference between children with DCD and those without, t_255_=-2.72, p=.007 – the 11 children with DCD had larger, earlier peak grip apertures, and made more additional acceleration on perturbed trials, as compared to children without DCD.

By contrast to the MABC-2, children with low versus high parent ratings of movement coordination (DCDQ’07) did not differ significantly in their factor scores, although the direction of effects was equivalent to those in the MABC-2 groups. Finally, while responding to reviewers’ comments, we discovered several significant differences in the factor scores for participants who performed the task in the dark versus in the light, who used their dominant versus their non-dominant hand to reach, and based on their gender. Analysis of these categorical variables, along with age and other participant-specific predictors, is beyond the current scope and will be dealt with fully elsewhere (Blanchard et al., in preparation).

## Discussion

The aim of this study was to investigate which kinematic variables of the visual online control of reaching and grasping movements could predict the standard scores of MABC-2 and DCDQ’07. Our results show that two factors extracted from a large number of movement variables provided strong predictions of MABC-2 performance, most strongly for aiming and catching scores. None of the factors individually or combined significantly predicted the DCDQ’07 scores. In the following, we discuss the relationships between reaching and grasping and the MABC-2, focussing on the measurement and analysis of movements in double-step perturbation tasks.

### Sensorimotor processes underlying reaching, grasping, catching, and aiming

Performance of our reaching and grasping task requires accurate planning, generation, and visual online control of reach-to-grasp movements, including the coordination of reaching and grasping phases. From our results, the strongest predictor of MABC-2 scores (especially the aiming and catching component) was factor 6, which loaded heavily on measures of the additional acceleration and jerk on perturbed compared to unperturbed trials, movement path, curvature, the latency to initiate movement corrections, and peak grip aperture. This factor may represent the key sensorimotor processes required in visual online control. Following a change in target location, the ideal movement correction would comprise a change of direction towards the new target, but without an overall increase in movement speed (i.e., no additional acceleration or jerk), and with a minimum overall increase in movement path length and duration. Efficient corrections will thus have lower overall jerk, path, movement time, curvature, and correction latency. Factors 6 and 15 also loaded on the variability and relative timing of peak grip aperture. An ideal correction to the reaching component of the movement should not also require a correction to the grasping component. Children who correct their reaching movement optimally would not need to adjust their grasping movement - the time to peak grasp aperture could stay relatively constant relative to overall movement time. By contrast, children who fail to adjust their reaching movement efficiently might close their grasp onto the central target, then require an additional opening of the grasp for the peripheral target. On some trials, the initial grasp will be detected as the peak grip aperture, and on other trials peak aperture will occur on the second grasp. This double-grasping movement leads to greater variability in the measured relative time of peak grip aperture. Our result echoes an earlier finding in which children with DCD showed a much greater variability in grasp timing than typically developing children (Astill and Utley, 2008). The authors of this previous study suggested that children with coordination disorders may use a decomposition strategy to simplify the control of transport and grasp phases of catching by uncoupling these movement components.

While aiming and catching scores were best-predicted by the reaching and grasping factors (16% of variance in the MABC-2 explained), manual dexterity, and to a lesser extent balance scores, were also significantly predicted by reaching and grasping, with 10% and 4% of variance explained, respectively. Because scores across the three components of the MABC-2 are correlated (across 225 of our participants, manual dexterity component scores correlate with aiming and catching, r_223_=.342, and balance, r_223_=.525; aiming and catching correlates with balance, r_225_=.389), factors which predict one of the components are also likely to predict the others. This is likely due to general movement coordination ability, to general cognitive, attentional, or motivational factors which are common to the movement tasks, or to the fact that accurate control of the hands and arms also requires postural and balance control, leading to functional links in development of these abilities (Flatters et al.,2014).

The relationship between kinematic factors and the aiming and catching component of the MABC-2 (16% variance explained) was modest, given that, for example, the manual dexterity and balance components shared 28% of variance in our dataset. Nevertheless, we found no significant relationships at all between any of our kinematic factors and the DCDQ’07 parent questionnaire. This negative finding suggests that parents’ evaluations of how their child’s movement coordination ability compares with others’ should be interpreted cautiously. The DCDQ’07 alone may be unlikely to measure movement coordination skill, at least for reaching, grasping, aiming, and catching skills, although note that we did not measure the DCDQ’07 and the MABC-2 in the same children.

Finally, no significant relationship was found between reaction time variables or the factors that loaded heavily on them, and the MABC-2 scores. Henderson and colleagues (1992) observed both prolonged simple reaction time and movement time in simple aiming in children with DCD. However, Hyde and Wilson, (2013) found that RT in children with mild to moderate motor impairments (DCD) was not significantly different than in TD children. The authors used this result as evidence to support the claim that there is not a basic information processing impairment in children with DCD. However, earlier work (Henderson et al., 1992; Hyde and Wilson, 2011a, 2011b) found longer RT to targets in children with motor impairment compared to matched controls, and is also consistent with other literature showing longer RT to external stimuli in children with DCD under lighting conditions which did not permit children to see their moving limb (Wilson and Hyde, 2013; Wilson and McKenzie, 1998). It is likely, then, that differences in RT between groups of children with and without DCD or other motor disorders are task-dependent (Mon-Williams et al., 2005).

### Measuring rapid visual online control

One important aspect of the present work concerns the method of measuring online movement corrections. Many different methods are possible and useful in different contexts (Holmes and Dakwar, 2015; Oostwoud Wijdenes et al., 2014), but the optimal methods in the present context involve fitting a model to the expected velocity or acceleration curves that arise from correction movements (Veerman et al., 2008). Previous studies have used manual estimation of trajectory deviations (e.g., Hyde and Wilson, 2011b, 2013), but these methods are time-consuming and prone to experimenter bias or error. Our fully-automated analysis extracted a number of variables reflecting the latency, velocity, and acceleration of movement corrections, and performed this analysis on both individual trials and the averaged velocities across trials. We found important discrepancies between methods of measuring online control: First, using average movement trajectories can result in substantially shorter correction latencies than using individual trial-by-trial analyses, due to temporal smoothing and broadening of velocity curves. The mean correction latency based on individual trials was 342 ms, while the mean based on the means of trials was 299 ms. Second, correction latencies based on differences in total movement time, which confounds correction latency with the post-correction movement time, were just 235 ms, more than 100 ms less than that of the individual trial-by-trial analysis. The additional movement time following a movement correction will be lower in children who reach faster or straighter overall, or who execute a faster correction movement. Indeed, the 107 ms difference between our preferred measure of correction latency and the additional movement time suggests that children increase their movement speed substantially after the target change, 'catching-up'. While our preferred correction latency measure was longest for the lower target location, the additional movement time required was longest for the upper target location. Moving the arm upwards probably requires more effort than moving downwards, so the post-correction movement direction may well influence overall movement time. Correction latencies from different studies can only meaningfully be compared if identical methods were used to measure them.

We chose to extract as many variables as possible from the reaching and grasping movements in an attempt to fully capture the differences in movement between children and conditions. With 87 extracted variables, the problem of multiple comparisons and collinearity arises, which we addressed by reducing the data to 17 relatively independent factors (cf Naish et al., 2013). An alternative, preferable, but computationally-expensive approach is to fit a series of low-dimensional models to the raw velocity data, and to analyse only the model parameters across participants and conditions. This analysis of ‘sub-movements’ is based on the minimum-jerk model, and may account well for online movement corrections (Flash and Henis, 1991). This approach, using constrained non-linear optimisation in Matlab, was investigated for analysis of the current dataset. However, with 262 participants and 40 trials, the computer processor time alone was likely to take several months! We will use this technique for future work.

### Continuous versus discrete groups of movement ability

Our approach to data analysis was continuous, in that we did not set out to create two distinct groups consisting of children with DCD and TD children. Rather, we explored motor abilities across the spectrum, eliminating the difficulties that arise when trying to categorise DCD, which is well known for its heterogeneity (Zwicker et al., 2012). We have noted that diagnosis of DCD is incomplete in the local population, and variable between groups of children, for example from different schools or administrative areas. Furthermore, we found that some children with a diagnosis of DCD performed perfectly well on the MABC-2. This could be due either to the wrong diagnosis being made, an intervention having been effective, developmental improvements since diagnosis, or to the inadequacy of the MABC-2 as a diagnostic instrument (Venetsanou et al., 2011). In the absence of a diagnosis, then, any division of continuous MABC-2 data into discrete clinical (i.e., ≤ 5^th^ percentile) or pre-clinical (i.e. 6^th^ < percentile ≤ 16^th^) categories is arbitrary, and, we suggest, likely to obscure the underlying, probably continuous, relationships between individual movement parameters and performance on the MABC-2. We have also noted that ceiling effects and the non-parametric distribution of data in some MABC-2 tasks limits the sensitivity of the MABC-2 to measure the full, continuous range of movement skill, and may have substantial implications for interpreting the standard cut-offs at the 5^th^ and 16^th^ percentiles (French et al., in preparation).

## Conclusions

Our results support the interpretation that impaired visual online control is a strong predictor of performance on standard tests of movement ability, as are often used to diagnose developmental movement disorders. The visual online control task developed for this study provides a continuous and high-resolution measurement, and is directly comparable between adults and children, which makes it a promising task for further study. The present results show that children who are poor at aiming and catching are also particularly poor at the online control of reaching and grasping.

## Acknowledgements

Our work was funded by the Medical Research Council (grant number MR/K014250/1 to NPH).

## References

American Psychiatric Association (2000). Diagnostic and Statistical Manual of Mental Disorders, Fourth Edition: DSM-IV-TR®. American Psychiatric Association.

American Psychiatric Association (2013). Diagnostic and Statistical Manual of Mental Disorders. Fifth Edition. American Psychiatric Association Available at: http://psychiatryonline.org/doi/book/10.1176/appi.books.9780890425596 [Accessed January 19, 2016].

Astill, S., and Utley, A. (2008). Coupling of the reach and grasp phase during catching in children with developmental coordination disorder. J. Mot. Behav. 40, 315–323. doi:10.3200/JMBR.40.4.315-324.

Blanchard, C.C.V., McGlashan, H. L., French, B., and Holmes, N. P. (in preparation). Development of online control.

Blank, R., Smits-Engelsman, B., Polatajko, H., and Wilson, P. (2012). European Academy for Childhood Disability (EACD): Recommendations on the definition, diagnosis and intervention of developmental coordination disorder (long version)*. Dev. Med. Child Neurol. 54, 54–93. doi:10.1111/j.1469-8749.2011.04171.x.

Cameron, B. D., Cheng, D. T., Chua, R., van Donkelaar, P., and Binsted, G. (2013). Explicit knowledge and real-time action control: anticipating a change does not make us respond more quickly. Exp. Brain Res. 229, 359–372. doi:10.1007/s00221-013-3401-z.

Castiello, U., Paulignan, Y., and Jeannerod, M. (1991). Temporal dissociation of motor responses and subjective awareness. A study in normal subjects. Brain J. Neurol. 114 (Pt 6), 2639–2655.

Comrey, A. L., and Lee, H. B. (1992). A first course in factor analysis. 2nd Edition. Lawrence Earlbaum, Hove.

Conners, C. K. (2008). Conners (3rd ed.). Pearson scientific.

Desmurget, M., and Grafton, S. (2000). Forward modeling allows feedback control for fast reaching movements. Trends Cogn. Sci. 4, 423–431.

Dunn, L., Dunn, L., Whetton, C., and Burley, J. (1997). British Picture Vocabulary Scale. 2nd edition. Windsor, Berks: NFER-Nelson.

Elliot, C. D. (1996). British Ability Scales II. Windsor, England: NFER-Nelson.

Elze, T., and Tanner, T. G. (2012). Temporal Properties of Liquid Crystal Displays: Implications for Vision Science Experiments. PLoS ONE 7. doi:10.1371/journal.pone.0044048.

Farnè, A., Roy, A. C., Paulignan, Y., Rode, G., Rossetti, Y., Boisson, D., et al. (2003). Visuo-motor control of the ipsilateral hand: evidence from right brain-damaged patients. Neuropsychologia 41, 739–757. doi:10.1016/S0028-3932(02)00177-X.

Flash, T., and Henis, E. (1991). Arm trajectory modifications during reaching towards visual targets. J. Cogn. Neurosci. 3, 220–230. doi:10.1162/jocn.1991.3.3.220.

Flatters, I., Mushtaq, F., Hill, L. J. B., Holt, R. J., Wilkie, R. M., and Mon-Williams, M. A. (2014). The relationship between a child’s postural stability and manual dexterity. Exp. Brain Res. 232, 2907–2917. doi:10.1007/s00221-014-3947-4.

Flora, D. B., LaBrish, C., and Chalmers, R. P. (2012). Old and New Ideas for Data Screening and Assumption Testing for Exploratory and Confirmatory Factor Analysis. Front. Psychol. 3. doi:10.3389/fpsyg.2012.00055.

French, B., Blanchard, C., McGlashan, H. L., and Holmes, N. P. (in preparation). Towards non-parametric, bootstrapped norms for the Movement Assessment Battery for Children-2 (MABC-2).

Goodale, M. A., Pelisson, D., and Prablanc, C. (1986). Large adjustments in visually guided reaching do not depend on vision of the hand or perception of target displacement. Nature 320, 748–750. doi:10.1038/320748a0.

Henderson, L., Rose, P., and Henderson, S. (1992). Reaction Time and Movement Time in Children with a Developmental Coordination Disorder. J. Child Psychol. Psychiatry 33, 895–905. doi: 10.1111/j.1469-7610.1992.tb01963.x.

Henderson, S.E., Sugden, D.A, and Barnett, A.L (2007). Movement Assessment Battery for Children-2. Second Edition (Movement ABC-2). Examiner’s manual. London: Harcourt Assessment.

Hill, L. J. B., Mushtaq, F., O’Neill, L., Flatters, I., Williams, J. H. G., & Mon-Williams, M. (2016). The relationship between manual coordination and mental health. European Child & Adolescent Psychiatry, 25, 283–295. doi:10.1007/s00787-015-0732-2.

Holmes, N. P., and Dakwar, A. R. (2015). Online control of reaching and pointing to visual, auditory, and multimodal targets: Effects of target modality and method of determining correction latency. Vision Res. 117, 105–116. doi:10.1016/j.visres.2015.08.019.

Holt, R. J., Lefevre, A. S., Flatters, I. J., Culmer, P. R., Wilkie, R. M., Henson, B. W., Bingham, G. P., and Mon-Williams, M. A. (2013). Grasping the changes seen in older adults when reaching for objects of varied texture. PLoS ONE, 8(7), e69040. doi:10.1371/journal.pone.0069040.

Hyde, C. E., and Wilson, P. H. (2013). Impaired online control in children with developmental coordination disorder reflects developmental immaturity. Dev. Neuropsychol. 38, 81–97. doi:10.1080/87565641.2012.718820.

Hyde, C., and Wilson, P. (2011a). Online motor control in children with developmental coordination disorder: chronometric analysis of double-step reaching performance. Child Care Health Dev. 37, 111–122. doi:10.1111/j.1365-2214.2010.01131.x.

Hyde, C., and Wilson, P. H. (2011b). Dissecting online control in Developmental Coordination Disorder: a kinematic analysis of double-step reaching. Brain Cogn. 75, 232–241. doi:10.1016/j.bandc.2010.12.004.

Mon-Williams, M. A., Tresilian, J. R., Bell, V. E., Coppard, V. L., Nixdorf, M., and Carson, R. G. (2005) The preparation of reach-to-grasp movements in adults, children, and children with movement problems. Q. J. Exp. Psychol., 58, 1249–1263. doi:10.1080/02724980443000575.

Naish, K. R., Reader, A. T., Houston-Price, C., Bremner, A. J., and Holmes, N. P. (2013). To eat or not to eat? Kinematics and muscle activity of reach-to-grasp movements are influenced by the action goal, but observers do not detect these differences. Exp. Brain Res. 225, 261–275.

Oostwoud Wijdenes, L., Brenner, E., and Smeets, J. B. J. (2014). Analysis of methods to determine the latency of online movement adjustments. Behav. Res. Methods 46, 131–139. doi:10.3758/s13428-013-0349-7.

Paulignan, Y., Jeannerod, M., MacKenzie, C., and Marteniuk, R. (1991a). Selective perturbation of visual input during prehension movements. 2. The effects of changing object size. Exp. Brain Res. 87, 407–420.

Paulignan, Y., MacKenzie, C., Marteniuk, R., and Jeannerod, M. (1991b). Selective perturbation of visual input during prehension movements. 1. The effects of changing object position. Exp. Brain Res. 83, 502–512.

Plumb, M. S., Wilson, A. D., Mulroue, A., Brockman, A., Williams, J. H. G., and Mon-Williams, M. (2008). Online corrections in children with and without DCD. Hum. Mov. Sci. 27, 695–704. doi:10.1016/j.humov.2007.11.004.

Prablanc, C., and Martin, O. (1992). Automatic control during hand reaching at undetected two-dimensional target displacements. J. Neurophysiol. 67, 455–469.

Raethjen, J., Pawlas, F., Lindemann, M., Wenzelburger, R., and Deuschl, G. (2000). Determinants of physiologic tremor in a large normal population. Clin. Neurophysiol. 111, 1825–1837.

Ruddock, S., Piek, J., Sugden, D., Morris, S., Hyde, C., Caeyenberghs, K., et al. (2014). Coupling online control and inhibitory systems in children with Developmental Coordination Disorder: Goal-directed reaching. Res. Dev. Disabil. 36C, 244–255. doi:10.1016/j.ridd.2014.10.013.

Swanson, J., Deutsch, C., Cantwell, D., Posner, M., Kennedy, J., Barr, C., et al. (2001). Genes and attention-deficit hyperactivity disorder. Clin. Neurosci. Res., 207–216.

Swanson, J. M., Schuck, S., Porter, M. M., Carlson, C., Hartman, C. A., Sergeant, J. A., et al. (2012). Categorical and Dimensional Definitions and Evaluations of Symptoms of ADHD: History of the SNAP and the SWAN Rating Scales. Int. J. Educ. Psychol. Assess. 10, 51–70.

Van Braeckel, K., Butcher, P. R., Geuze, R. H., Stremmelaar, E. F., and Bouma, A. (2007). Movement adaptations in 7- to 10-year-old typically developing children: evidence for a transition in feedback-based motor control. Hum. Mov. Sci. 26, 927–942. doi:10.1016/j.humov.2007.07.010.

Veerman, M. M., Brenner, E., and Smeets, J. B. J. (2008). The latency for correcting a movement depends on the visual attribute that defines the target. Exp. Brain Res. 187, 219–228. doi:10.1007/s00221-008-1296-x.

Venetsanou, F., Kambas, A., Ellinoudis, T., Fatouros, I., Giannakidou, D., and Kourtessis, T. (2011). Can the movement assessment battery for children-test be the “gold standard” for the motor assessment of children with Developmental Coordination Disorder? Res. Dev. Disabil. 32, 1–10. doi:10.1016/j.ridd.2010.09.006.

Wilson, B. N., Crawford, S. G., Green, D., Roberts, G., Aylott, A., and Kaplan, B. J. (2009). Psychometric Properties of the Revised Developmental Coordination Disorder Questionnaire. Phys. Occup. Ther. Pediatr. 29, 182–202. doi:10.1080/01942630902784761.

Wilson, B. N., Kaplan, B. J., Crawford, S. G., Campbell, A., and Dewey, D. (2000). Reliability and Validity of a Parent Questionnaire on Childhood Motor Skills. Am. J. Occup. Ther. 54, 484–493. doi:10.5014/ajot.54.5.484.

Wilson, P. H., and Hyde, C. (2013). The development of rapid online control in children aged 6-12 years: reaching performance. Hum. Mov. Sci. 32, 1138–1150. doi:10.1016/j.humov.2013.02.008.

Wilson, P. H., and McKenzie, B. E. (1998). Information Processing Deficits Associated with Developmental Coordination Disorder: A Meta-analysis of Research Findings. J. Child Psychol. Psychiatry 39, 829–840. doi:10.1111/1469-7610.00384.

Wilson, P. H., Ruddock, S., Smits-Engelsman, B., Polatajko, H., and Blank, R. (2013). Understanding performance deficits in developmental coordination disorder: a meta-analysis of recent research. Dev. Med. Child Neurol. 55, 217–228. doi:10.1111/j.1469- 8749.2012.04436.x.

Wolpert, D. M., Miall, R. C., and Kawato, M. (1998). Internal models in the cerebellum. Trends Cogn. Sci. 2, 338–347.

Zwicker, J. G., Missiuna, C., Harris, S. R., and Boyd, L. A. (2012). Developmental coordination disorder: a review and update. Eur. J. Paediatr. Neurol. 16, 573–581. doi:10.1016/j.ejpn.2012.05.005.

